# Evolutionary dynamics of abundant 7 bp satellites in the genome of *Drosophila virilis*

**DOI:** 10.1101/693077

**Authors:** Jullien M. Flynn, Manyuan Long, Rod A. Wing, Andrew G. Clark

## Abstract

The factors that drive the rapid changes in satellite DNA genomic composition we see in eukaryotes are not well understood. *Drosophila virilis* has one of the highest relative amounts of simple satellites of any organism that has been studied, with an estimated >40% of its genome composed of a few related 7 bp satellites. Here we use *D. virilis* as a model to understand technical biases affecting satellite sequencing and the evolutionary processes that drive satellite composition. By analyzing sequencing data from Illumina, PacBio, and Nanopore platforms, we identify platform-specific biases and suggest best practices for accurate characterization of satellites by sequencing. We use comparative genomics and cytogenetics to demonstrate that the highly abundant satellite family arose from a related satellite in the branch leading to the virilis phylad 4.5 - 11 million years ago before exploding in abundance in some species of the clade. The most abundant satellite is conserved in sequence and location in the pericentromeric region but has diverged widely in abundance among species, whereas the satellites nearest the centromere are rapidly turning over in sequence composition. By analyzing multiple strains of *D. virilis*, we saw that one centromere-proximal satellite is increasing in abundance along a geographical gradient while the other is contracting in an anti-correlated manner, suggesting ongoing conflicts at the centromere. In conclusion, we illuminate several key attributes of satellite evolutionary dynamics that we hypothesize to be driven by processes like selection, meiotic drive, and constraints on satellite sequence and abundance.

## Introduction

Repetitive DNA is abundant in most eukaryotic genomes, and is now understood to be correlated with the manifold variation in genome size across the tree of life (Elliott and Gregory 2015). For most species, transposable elements (TEs) dominate the repeat landscape, including in humans, plants, and *Drosophila melanogaster*. Satellite DNA, which is characterized by tandem repeats spanning long arrays, very rarely has dominated a genome to a similar extent as TEs. An unprecedented case is that of *Drosophila virilis*, the Drosophila species with the largest estimated genome size (up to 389 Mb) (Bosco *et al.* 2007), where some 40% of the genome is comprised of just three simple 7-mer satellites: AAACTAC, AAACTAT, and AAATTAC (Gall *et al.* 1971; Gall and Atherton 1974). Since the 1970s, there has been no follow-up to validate the amount of 7-mers with modern techniques, or evolutionary studies to understand how and why these satellite repeats expanded so explosively. The genomic composition of simple satellites in *D. virilis* provides an excellent model for an investigation of the evolutionary dynamics involved in their expansion in the genome as well as the technical challenges facing simple satellite analysis.

Satellites are rapidly evolving in sequence and copy number, and there is a high level of variation in satellite content among and within species (Wei *et al.* 2014, 2018). The reasons for such dramatic variation is not well understood, and cannot be fully explained by current models. Satellites have been long hypothesized to be slightly deleterious and therefore governed primarily by the strength of negative selection (Ohno 1972). However, the amount of satellite in the genome that causes negative effects that could be selected against depends on many factors and cannot be easily predicted (Charlesworth *et al.* 1994; Gregory 2001). The fact that most organisms have satellite repeats in or near centromeres suggests that they are important for centromere function. Satellite repeats can also be important for maintenance of the chromocenter and packaging of chromosomes in the nucleus (Jagannathan *et al.* 2018, 2019), and the transcripts of some satellites may be essential for fertility (Mills *et al*. 2019). In heterozygotes with alleles that differ in pericentromeric satellite sequence or abundance, one allele may assemble a stronger kinetochore during female meiosis I, increasing its probability of transmission into the egg (rather than polar bodies). This transmission advantage, known as centromere drive, allows satellites to rapidly change in composition in the population, regardless of their whole-organism fitness effects (Henikoff et al. 2001). If satellite DNA is an essential component of genomes or is only a burden (i.e. is selfish), it is still not clear why some species have almost no pericentromeric satellite DNA while others, like *D. virilis*, possess pericentromeric satellites that make up almost half of the genome.

Comparing the satellites of *D. virilis* to those of its sister species can elucidate when the abundant satellites arose, and how rapidly their copy numbers and sequences evolved. *D. virilis* is 4.5 MY diverged from its sister species *D. novamexicana* and *D. americana*, which are both restricted to North America, unlike globally-distributed *D. virilis* (Caletka and McAllister 2004). *D. novamexicana* and *D. americana* have a smaller estimated genome size than *D. virilis* (∼250 Mb vs. 389 Mb), suggesting these species may have less satellite content (Bosco *et al.* 2007). Additionally, using intra-species comparisons across global populations can give indications about factors that may be influencing satellite dynamics. For example, in *D. melanogaster*, patterns of abundance of the *Prodsat* satellite closely mirror the migration patterns of species, suggesting an ongoing expansion of this satellite (Wei *et al.* 2014). Genetic drift or meiotic drive may contribute to patterns of geographical gradients of satellite abundance. We can also use intra-species data to pose hypotheses about non-neutral processes that may be driving satellite content. Previous work has shown evidence for conflicts or trade-offs between satellites within the genome, and these constraints can be illuminated by analyzing satellites in several strains (Flynn *et al.* 2017, 2018).

Genome-wide characterization of satellites has taken off since high-throughput sequencing has become widely available. We have learned from several informative studies about the sequences and relative abundances of satellites in various species (Pavlek *et al.* 2015; Flynn *et al.* 2017; de Lima *et al.* 2017; Wei *et al.* 2018), but technical challenges may prevent accurate quantitative estimates. Satellites may be more prone to errors or biases in the sequencing process that do not affect the better studied regions of the genome. Satellites are difficult to assemble even with long-read sequencing (Chang and Larracuente 2019). The genome assembly of *D. virilis* is approximately half its estimated genome size by flow cytometry (∼200 Mb vs 389 Mb) (Bosco *et al.* 2007), and it is likely that much of what is missing is simple satellite DNA. However, even using alignment-free raw read methods have not produced satellite DNA estimates that approach the amount that is missing from the genome assembly and was estimated from early work (Gall *et al.* 1971; Gall and Atherton 1974; Wei *et al.* 2018). Now, as long read sequencing is also being exploited to study satellites, we must evaluate satellite DNA abundance estimates to assess if there are platform-specific biases that may affect evolutionary analysis of satellite DNA.

The purpose of this paper is two-fold; first to explore the technical biases preventing accurate characterization and quantification of simple satellites, and second to use a comparative approach to understand the evolutionary dynamics of the extremely abundant 7mers in the *D. virilis* group. First, we characterize satellites in *D. virilis* sequencing data from different platforms and assess biases that affect accurate satellite characterization. We then use comparative genomics and cytogenetics in *D. virilis* and its sister species to understand the composition and changes in the highly abundant simple satellites. Finally we sequence multiple strains of *D. virilis* and sister species to estimate polymorphism in satellite abundance and infer processes that may be influencing their evolution. From this we infer that there are likely a variety of understudied processes affecting satellite DNA in this organism, including positive selection, meiotic drive, and constraints and trade-offs between satellites.

## RESULTS

### Technical biases in characterizing simple satellites from sequencing

#### Long-read genome assemblies have an under-representation of simple satellites

Long-read sequencing technologies have an advantage because of their long reads, but a disadvantage due to their high error rate, prompting a need for extensive alignments for error-correction and assembly. First we asked whether assemblies from long read technologies can better assemble simple satellite reads than the previous Sanger assembly. We compared the amount of simple 7-mer satellites (AAACTAC, AAACTAT, AAATTAC, AAACAAC) in three *D. virilis* genome assemblies: the CAF1 assembly produced from Sanger sequencing (Drosophila 12 Genomes Consortium *et al.* 2007), a PacBio assembly produced by our group by ∼100x coverage (available at https://www.ncbi.nlm.nih.gov/bioproject/?term=txid7214[Organism:noexp]), and a Nanopore assembly produced from ∼20x sequencing coverage (Miller *et al.* 2018). All assemblies were approximately the same size at ∼200 Mb. The PacBio and Nanopore assemblies contained a similarly low amount of simple 7-mer satellites, 29 and 28 kb, respectively. The CAF1 assembly, however contained 7.36 Mb of these satellites. This discrepancy is likely largely due to the difference in assembly algorithms used for short read and long read data. Long reads must be aligned and corrected to be incorporated into the assembly because of their high error rate, whereas this is not necessary for Sanger-based assemblies. Use of modified methods can improve assemblies of repetitive regions (Chang and Larracuente 2019), but for highly homogeneous simple satellites, whose arrays span 10-100x longer than the current maximum read length, it is practically impossible to produce a continuous assembly.

#### Simulations to assess simple repeat quantification from long read sequencing data

Due to assembly issues of simple satellites, they must be quantified from raw unassembled reads. Long read sequencing data poses a significant challenge because of the high error rate including a high indel rate in the raw reads. We therefore used two different approaches along with simulations to assess their accuracy. The first approach used k-Seek (Wei et al. 2014) to select repeat-rich reads and then Phobos (https://www.ruhr-uni-bochum.de/ecoevo/cm/cm_phobos.htm) to quantify satellites. This approach allows for *de novo* discovery of satellite sequences. We used Noise-Cancelling Repeat Finder (NCRF, Harris *et al.* 2019) for our second approach, providing our target satellites. Both methods are relatively sensitive to imperfect repeats, which we expect with the high error rate of long-read sequencing.

To evaluate our approaches, we created a mock *D. virilis*-like genome containing pericentromeric and centromeric repeats on each of five chromosomes (See Materials and Methods). We then simulated 10x PacBio reads from this genome, and then quantified satellites using both approaches. NCRF works by doing alignments of target satellites to the reads and allowing up to a user-specified maximum divergence. To determine the most appropriate maximum divergence, we simulated a range of values for this parameter from 18-30% and chose the lowest asymptotic value - which was 25% in this case (Figure S1). NCRF found almost the same amount of satellites that truly existed in the mock genome whereas the k-Seek + Phobos method only found about 20% (Figure 1A).

**Fig 1:**
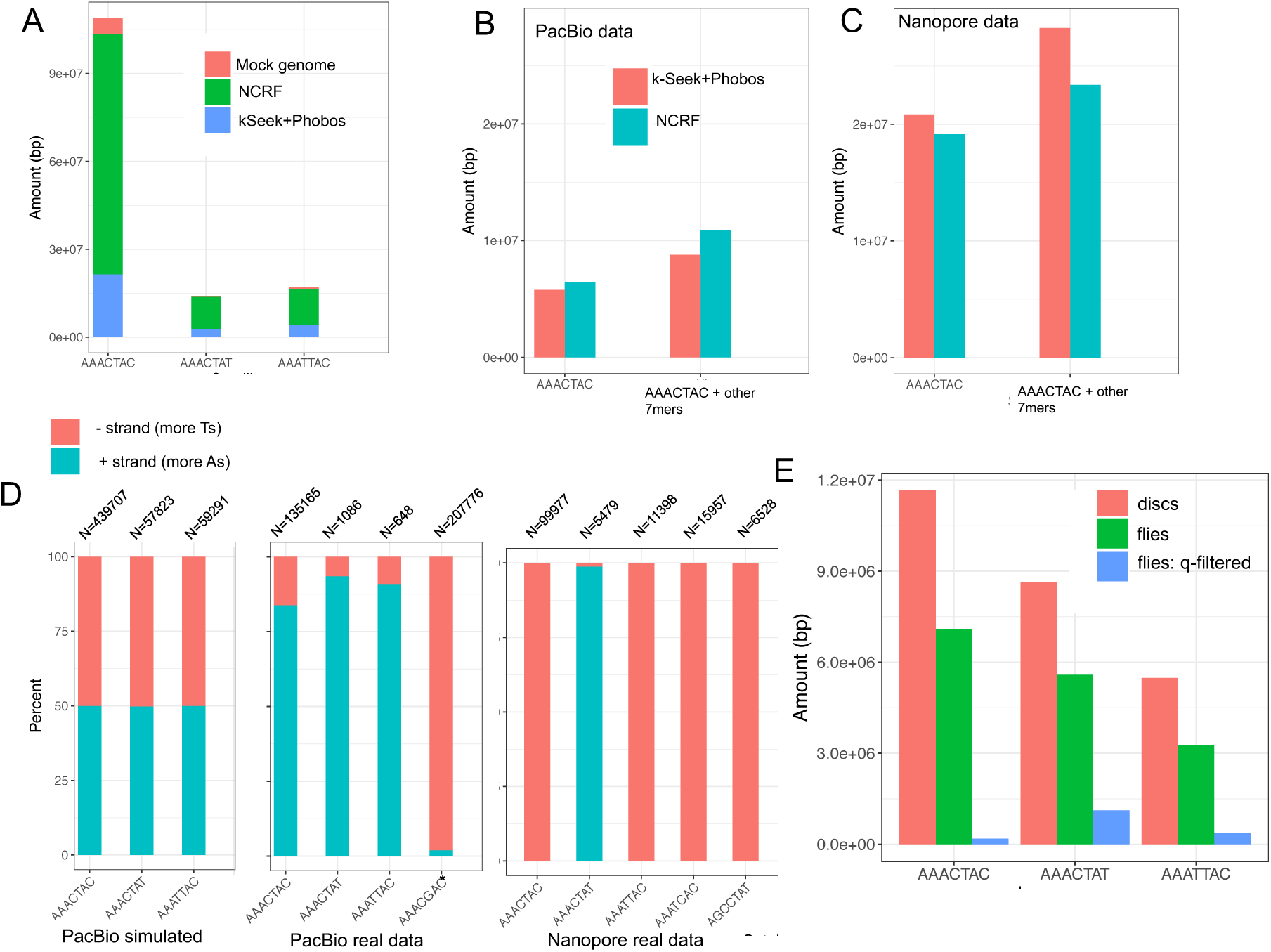
Issues in quantifying simple satellites in sequencing data (all data shown is *D. virilis*). (**A**) Cumulative stacked barplot comparing the performance of the two tested approaches on PacBio data simulated with PBSim from a mock genome. (**B)** Comparing the results of the two approaches on the real PacBio data; “other” refers to additional satellites in the family, including suspected artefactual ones (AAAGCAC for PacBio and AAATCAC + AGCCTAT for Nanopore). (**C**) Same as B for Nanopore data. (**D**) Strand biases in the sequenced satellites in long read sequencing data. Satellites with asterisks are suspected artefactual ones. N refers to the number of satellite regions of reads used for the calculation. (E) Amount of satellites quantified in different datasets: imaginal discs (pure diploid), compared to flies (some polyteny), and fly data that has been quality filtered.

#### The amounts and biases in simple 7mer repeats differ between Nanopore and PacBio sequencing reads

Next, we quantified simple satellites in the long-read data generated from our 100x PacBio sequencing and 20x Nanopore sequencing using the two approaches mentioned above. Unlike in the simulations, both approaches produced very similar (but lower than expected) estimates at 8.8-10.9 Mb for the PacBio data (Figure 1B). The Nanopore data contained almost 3 times the 7mer satellites compared to PacBio, with 23.4 - 28.2 Mb (Figure 1C). This may represent a platform-specific difference in the ability to sequence long arrays of simple tandem repeats. Both the PacBio reads and the Nanopore reads contained a greater amount of simple satellites than data produced in our lab previously with Illumina HiSeq sequencing (Wei et al. 2018), however did not approach the estimated >100 Mb in the genome.

Both the PacBio and Nanopore reads contained large amounts of what we expect to be artefactual repeats, which were found with the k-Seek + Phobos approach, and validated with NCRF. NCRF found 4.4 Mb (normalized to 1x genome coverage) of AAACGAC in the PacBio reads. This satellite was not found in the Nanopore data or Illumina data (this and previous studies) or in previous studies that characterized the most abundant satellites in *D. virilis*. Manual inspection proved that the AAACGAC satellite was the true consensus found in long arrays in the reads and did not represent an error in our approaches’ characterization of satellites. Similarly, AAATCAC, AGCCTAT, ACAGGCT, and AATGG were found in megabase quantities (after normalization) in the Nanopore data - whereas these satellites were not found in Illumina or PacBio data. We suggest these satellites are also technical artifacts introduced at the base-calling level.

In the PacBio data, the relative amounts of 7mer satellites (AAACTAC, AAACTAT, and AAATTAC) were lower than expected. This additional evidence led us to hypothesize that there were context-specific errors in our PacBio data affecting our particular satellites. If the sequencing were unbiased, we would expect to have an equal amount of satellites being detected on reads coming from both DNA strands. We evaluated the strand bias in the simulated and real long-read data for the three most abundant true satellites, as well as some artefactual satellites. We arbitrarily label the positive strand as AAACTAC and the negative strand as GTAGTTT, etc. In the simulated data, the positive and negative strands of satellites were detected in equal amounts (Figure 1D). However, there was a strong strand bias for all satellites in both the PacBio and Nanopore data (Figure 1D). For PacBio, the real satellites AAACTAC, AAACTAT, AAATTAC had a positive strand bias, whereas the artefactual satellite had a negative strand bias: 98% of the reads with this satellite were from the negative strand. Based on communication with PacBio representatives, this issue seemed to be caused by context-specific issues with base calling algorithms used for this sequencing run. As base calling algorithms improve, these issues will likely begin to be remedied. In fact, we received PacBio Circular Consensus Sequencing or “HiFi” data for a closely related species, *D. americana*, and the base-calling issue was remedied. In the Nanopore data, strand biases were even more extreme: the negative strand was sequenced almost exclusively for real satellites AAACTAC and AAATTAC and suspect satellite AAATCAC. However, the AAACTAT real satellite was sequenced almost exclusively on the positive strand. In this case, strand biases may be caused by unsequenceable secondary structures developing more frequently on one strand of the satellite DNA than the other. We analyzed Illumina NextSeq reads for *D. virilis*, and no such strand bias was found.

#### D. virilis whole-flies have 40% less pericentromeric satellites than non-polytene tissue

Polyteny occurs in all differentiated tissues of Dipterans, and is characterized by multiple rounds of local DNA replication within the same nucleus and without cell division, a process known as endoreduplication (Smith and Orr-Weaver 1991; Kim *et al.* 2011). However, the pericentromeric heterochromatin, where most satellite DNA is located, is under-replicated (Belyaeva *et al.* 1998). It has never been tested if the level of polyteny in an adult fly makes a difference in the estimate of satellites per genome. Thus, we sequenced adult male flies (which have multiple polytene tissues) and imaginal discs (which are diploid) from male larvae and compared the amount of simple satellites in these datasets. We used Illumina sequencing and PCR-free library preparations to reduce known PCR bias (Wei *et al.* 2018). We found that for each of the four most abundant 7mer satellites in the *D. virilis* genome, there was approximately 40% less in the flies compared to the imaginal discs (Figure 1E). This pattern is not observed for microsatellites which are known to localize outside of pericentromeric heterochromatin (Figure S2A). We also analyzed publicly available *D. melanogaster* data, including flies, imaginal discs, and salivary glands (which are the most extreme in polyteny), and observed this same pattern of under-replication of satellite repeats in polytene tissues (Figure S2B and S2C).

#### Reads with tandem repeats had lower quality scores in Illumina data

Upon inspection with FastQC of our data from the polyteny analysis, we found a bimodal distribution of quality scores, with one peak at 22 and another at 37 (Figure S3A). After filtering low quality reads, the majority of the reads with simple satellites were removed (Figure S3). The quantity of satellites was reduced by ∼15 x after quality filtering (Figure 1E). It is apparent that in our dataset, simple satellite-containing reads were highly enriched for low quality scores. We examined other published *D. virilis* Illumina datasets to evaluate if this issue existed in other sequencing runs. Two other datasets were available and the one that was produced on the Illumina NextSeq platform like our data (Miller *et al.* 2018) showed the same pattern of biased quality scores in repetitive reads (Figure S4). The dataset produced on the HiSeq platform (Wei *et al.* 2018) did not show this pattern. It should be noted however that the amounts of 7mer satellites sequenced in the NextSeq datasets were higher than the HiSeq dataset. Our libraries were multiplexed with other non-*D. virilis* group samples from unrelated projects and only represented ∼20% of the total sequenced lane so that we would not have issues related to low complexity. We also noticed this pattern (but less dramatically) in our Illumina sequencing of multiple strains.

### Related species have similar but fewer simple repeats

*D. novamexicana* and *americana* which are 0.38 MY diverged from each other, are sister species of *D. virilis*, which is approximately 4.5 MY diverged (Caletka and McAllister 2004) (Figure 2A). We sequenced these species with high coverage PacBio runs and characterized and quantified satellites. We emphasize the comparison of relative satellite amounts since all are likely under-represented. *D. americana* was sequenced with PacBio HiFi reads, which eliminated artefactual satellites, but make quantitative comparisons difficult since different chemistries have different efficiencies of sequencing satellites. Nevertheless, we also found a high enrichment of 7bp satellites in *D. novamexicana* and *D. americana* (Fig 2B). Interestingly, we found the most abundant satellite in *D. virilis*, AAACTAC, is also the most abundant in *D. novamexicana* and *D. americana*, albeit with about half the total amount. The second and third most abundant repeats, AAACTAT and AAATTAC, however were not present in long tandem arrays in *D. novamexicana*. The second most abundant satellite in *D. novamexicana* and *americana* was AAACAAC, whereas in *D. virilis* there is only a few kilobases.

**Fig 2.**
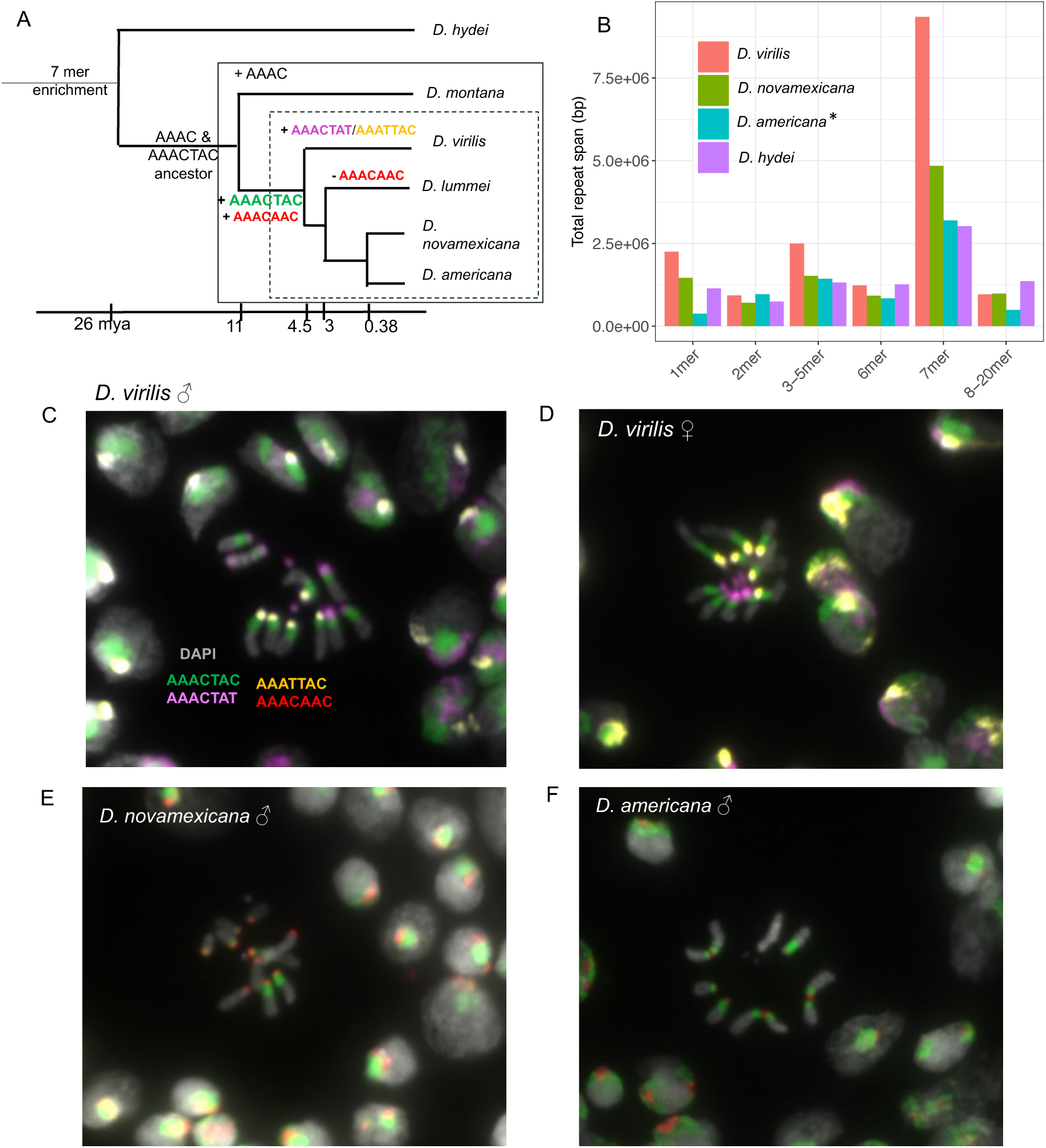
Comparative analysis of simple satellites in the *D. virilis* group. (A) Phylogeny demonstrating when satellites arose (+) and were lost (-). Dashed box: virilis phylad; solid box: virilis group. (B) Total amount of different unit lengths (k-mers) of satellites across four related species. The (*) for *D. americana* indicates that it was sequenced with PacBio HiFi reads, whereas the other species were sequenced with chemistry version 2.0. (C) DNA-FISH image of *D. virilis* male mitotic cells (D) DNA-FISH image of *D. virilis* female mitotic cells (E) DNA-FISH image of *D. novamexicana* male mitotic cells (F) DNA-FISH image of *D. americana*. Up to three different fluorescent probes were used each time.

By analyzing sequencing data in more diverged species, we can infer when the AAACTAC satellite family arose. *D. hydei* is approximately 26 MY diverged from *D. virilis* (Izumitani *et al.* 2016), and we had PacBio long read data for this species. Here 7 bp satellites are again the most enriched (Fig 2B), but the sequences are unrelated to those in *D. virilis* (ACCCATG, AAAGGTC from PacBio data). We analyzed Illumina data for *D. montana*, another member of the virilis group that is 7-11 MY diverged from *D. virilis* (Ostrega and Thompson 1986; Spicer and Bell 2002) (Figure 2A). This species does not have any AAACTAC family satellites, and in fact no enrichment of 7 bp satellites. The most abundant satellite in *D. montana* is AAAC. From these data, we infer that the AAACTAC family of satellites arose in the clade leading to the *D. virilis* phylad 4.5-11 MYA. We also analyzed Illumina sequencing data for *D. lummei*, which is 3 MY diverged from *D. novamexicana/americana* (Fig 2A). AAACTAC is conserved in *D. lummei,* but it is the only enriched 7 bp satellite in this species and its relative estimated abundance is lower than the other three *D. virilis* phylad species.

### Complex satellites are also abundant in *D. virilis* group genomes

We searched the high-quality genome assemblies for complex satellites (defined here as unit lengths greater than 20 bp). In *D. virilis,* we found a 36-bp satellite AAAACGACATAACTCCGCGCGGAGATATGACGTTCC making up ∼800 kb of the assembly. This satellite was found in previous studies and is thought to be associated with the possibly mobile element pDv (Zelentsova et al 1986, Heikkinen et al. 1995). In *D. novamexicana*, we found a 32 bp satellite AAAAGCTGATTGCTATATGTGCAATAGCTGAC along with a related 29 bp satellite. The 32 bp satellite spanned over 1.1 Mb on a single 3 Mb contig in the *D. novamexicana* assembly. The non-satellite portion of the contig had similarity to chromosome 6 (dot chromosome/Muller element F) (Figure S5). In *D. americana*, we found this identical 32 bp satellite, but in total its span was only ∼150 kb. In all *D. virilis* group species, we also found a series of similar satellites varying in size (150-500 bp) related to the previously described helitron central repeat that has expanded to tandem repeats in the virilis group (Dias *et al.* 2015).

### Fluorescence *in situ* hybridization reveals evolutionary dynamics of 7 bp repeats

The location of the 7 bp satellites on metaphase chromosomes has never been shown in the *D. virilis* group. From our sequencing data, we know that the AAACTAC satellite is conserved between *D. virilis, D. novamexicana,* and *D. americana*, but the abundance varies by approximately two-fold. The second most abundant satellites have turned over between *D. virilis* and *novamexicana*/*americana*. We used FISH of the most abundant 7mers (AAACTAC, AAACTAT, AAATTAC, AAACAAC) in these three sister species. *D. virilis* and *D. novamexicana* have the same karyotype with five acrocentric chromosomes plus the very small F element or “dot chromosome”. The strain of *D. americana* we used has centromere-centromere fusions between the X and 4^th^ chromosomes and the 2^nd^ and 3^rd^ chromosomes.

FISH results in *D. virilis* show that the most abundant satellite determined by sequencing, AAACTAC, is clearly the most abundant and occurs in approximately equal amounts in the pericentromeric region on the five pairs of large chromosomes. The Y chromosome appears to have slightly less AAACTAC satellite. The second and third most abundant satellites, AAATTAC and AAACTAT, are localized more proximally at or near the centromere. There are five single chromosomes having each of these satellite, indicating that one chromosome pair has different satellite content - which we hypothesized to be the X and Y. Based on differences between male and female FISH results (Figure 2C and 2D), we suggest the Y chromosome has AAACTAT at both distal ends of the chromosome and AAACTAC only flanking one end, whereas the X chromosome has the other centromeric repeat AAATTAC. We were also able to visualize the dot chromosomes in *D. virilis*, which we find is mostly composed of AAACTAT. The AAACAAC satellite is present in small amounts in *D. virilis*, very likely on a single chromosome (Figure S6).

We estimated that *D. novamexicana* has approximately half the AAACTAC as *D. virilis*, and visualizing it with FISH reveals a pattern that suggests aspects of its evolution. Its pericentromeric localization is conserved. One chromosome pair has the same amount of AAACTAC as *D. virilis*, whereas all other chromosomes have a very small amount (Figure 2E). Based on the FISH images, it appears that it is the 5th chromosome in *D. novamexicana* that has the greatest amount of pericentromeric AAACTAC conserved. The centromeric repeat on all major chromosomes is AAACAAC in *D. novamexicana* and *D. americana.* Our images illustrate clearly the centromere-centromere fusion between chromosome X-4 and 2-3 in *D. americana* with the satellites being maintained on both sides of the fusion (Figure 2F). None of the four simple satellite probes bound to the Y chromosome of *D. novamexicana* or *D. americana*. Based on the images we suggest that *D. americana* has an intermediate amount of pericentromeric AAACTAC satellite compared to *D. virilis* and *novamexicana*.

### Some satellite-containing reads are linked to TEs

We used RepeatMasker to detect if any of the reads containing satellites also contain transposable elements. TEs might be located in islands within the simple repeats at the centromere as in *D. melanogaster* (Chang *et al.* 2019) (Figure 3A), in more distal regions flanking the pericentromeric heterochromatin (Figure 3B), or some TEs may have inserted into long pericentromeric satellite arrays (Figure 3C). For AAACTAC (and its artefactual counterpart AAACGAC), ∼3.5% of reads (2473/75,364) also contained at least 500 bp of a TE insertion. In the satellite reads that also contain TE sequences, TEs were enriched at the beginning and ends of reads, concordant with the hypothesis that a high proportion of the TEs we found are flanking the long arrays of AAACTAC distally (Figure 3B). In order to understand how many reads would be expected to contain both satellites and TEs if TEs flanked this satellite and were not interspersed, we simulated a situation where a large satellite block flanked a large TE block. This simulation revealed a much smaller amount of reads containing both satellites and TEs (0.06 %). This result suggests that not only are TEs flanking the pericentromeric satellite AAACTAC, but there have likely been TE insertions into the satellite arrays. Likely flanking the proximal end of the pericentromeric satellite are the centromeric satellites AAACTAT or AAATTAC. 144 reads contained both AAACTAC and AAACTAT repeats (0.19% of AAACTAC reads) and 94 reads contained both AAACTAC and AAATTAC repeats (0.12% of AAACTAC reads). Based on our simulations, these proportions of overlapping reads are consistent with our expectation based on our FISH results that the pericentromeric and centromeric satellites have relatively clean boundaries and they directly flank each other.

**Fig 3.**
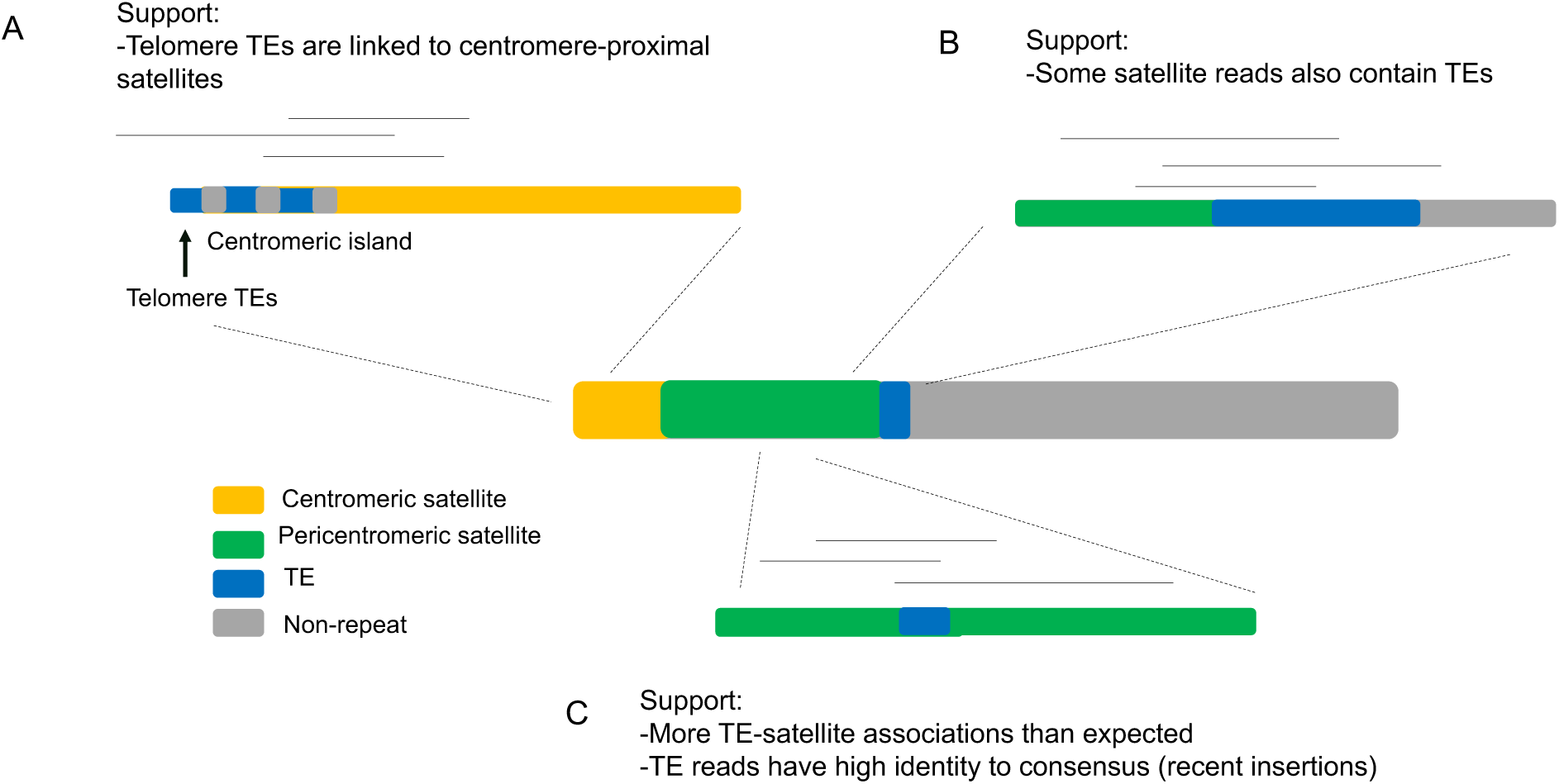
Transposable elements are in close proximity to satellite arrays. (**A**) Satellites near the centromere may be linked to potential islands of retroelements (Chang et al. 2019) and telomeric TEs. (**B**) Heterochromatic TEs flank satellite arrays. Histograms show the start and end position of transposable element in the satellite-rich read. (**C**) Transposable elements may have inserted into the satellite arrays.

Many of the TE insertions into satellites seemed to be very recent. 2080/2473 TE-containing AAACTAC reads had a TE insertion with less than 15% divergence from the Repbase consensus, which is the expected error rate of PacBio reads. These insertions included the superfamilies: DNA elements, LINE/CR1, LINE/I-Jockey, LINE/Penelope, LINE/R1, LTR/Copia, LTR/Gypsy, LTR/Pao, and Helitrons. We acknowledge however, that there may be a detection bias for insertions that are less divergent from the consensus. We also remind readers that there were likely fewer satellite reads sequenced than expected, and that this may have biased these results if satellite-only regions were sequenced less efficiently than satellite-TE regions. For centromeric satellites AAATTAC and AAACTAT, the results are more difficult to interpret since these were more strongly under-represented. However, 300/715 reads of AAACTAT and 49/385 reads AAATTAC contained TEs. Like the AAACTAC pericentromeric satellite, most TE insertions were low divergence from the Repbase consensus in the centromere-proximal satellite reads. 288/300 and 46/49 TE containing reads had a TE insertion > 500 bp with < 15% divergence for AAACTAT and AAATTAC, respectively.

There were differences in the TE composition of reads with different satellites. For the pericentromeric satellite AAACTAC, Gypsy-10_Dvi was the most enriched, followed by Helitrons (Helitron-1N1_DVir, Helitron-1_DVir, Helitron-2N1_DVir, Helitron-2_DVir). For the AAACTAT centromeric satellite, Gypsy-10_Dvi was again the most enriched, followed by Penelope. For the AAATTAC centromeric satellite, Gypsy-2_DVir was the most enriched followed by Penelope. In both AAACTAC and AAACTAT reads, CR1-1_DVi was the second or third most abundant TE. Interestingly, R1 was present in relatively high amounts in AAACTAC. In 110/132 of these R1-AAACTAC reads, rDNA sequences were not also linked. This suggests that some R1 elements, which are generally localized to rDNA loci, have jumped into or near satellite arrays. This is concordant with findings that some R1 elements are located outside rDNA loci in Drosophila (Stage and Eikbush 2009). All centromeres in *D. virilis* are acrocentric, meaning that the telomeric TEs Het-A and TART (Casacuberta and Pardue 2003) are likely near the centromere satellites. We found 12 reads linked to AAATTAC that contained matches to TART. Only two reads linked to any satellite contained a sequence matching HeT-A. We also used BLAST to detect matches between the genome assembly (masked from the 7mer satellites) and the 7mer satellite reads. We could not detect any unique regions of the genome that matched non-satellite sequence on the reads because they had low quality matches to hundreds of places in the genome each.

### Variation in *D. virilis* group global strains

*D. virilis* is globally distributed while its sister species are localized to North America, with *D. novamexicana* more restricted than *D. americana.* Patterns of variation in satellites may reveal potential mechanisms that can be hypothesized to be driving satellite evolution. Additionally, *D. americana* has a polymorphic fusion between the X and 4th chromosomes, so we may be able to identify differences in satellite composition associated with the fusion. This fusion has been shown to be currently undergoing meiotic drive, potentially mediated by a larger total centromere or pericentromere size in the fused strains compared to the non-fused strains (Stewart et al. 2019). On the other hand, chromosome fusions are often caused by Robertsonian translocations with loss of some non-essential DNA, which might include pericentromeric satellites (Schubert and Lysak 2011).

We used Illumina sequencing with PCR-free library preparation and k-Seek to estimate the abundance of 7mer satellites across 12 worldwide strains of *D. virilis*, eight strains of *D. americana* (including four strains that have the X-4 fusion and four that do not), and five strains of *D. novamexicana* (Table S1). All sequenced strains were male except a female of the *D. virilis* inbred strain 87 as a comparison. A PCA using only the four most abundant 7mers shows clustering of the three species, but the separation is much more dramatic in the PCA using the 20 most abundant simple satellites (Figure S7). Overall, *D. virilis* had the highest AAACTAC satellite content as well as the highest variation, with *D. americana* intermediate between *D. virilis* and *D. novamexicana* (Figure 4A). Using different normalization procedures including mapping and GC correction (see Materials and Methods), produced the same relative ranking of satellite abundances between species. In all cases, the inbred strain from which the genome sequence was produced had the lowest abundance of AAACTAC. In the case of *D. virilis*, this difference was very high. This was not due to a normalization bias as we did mapping-free normalization.

**Fig 4.**
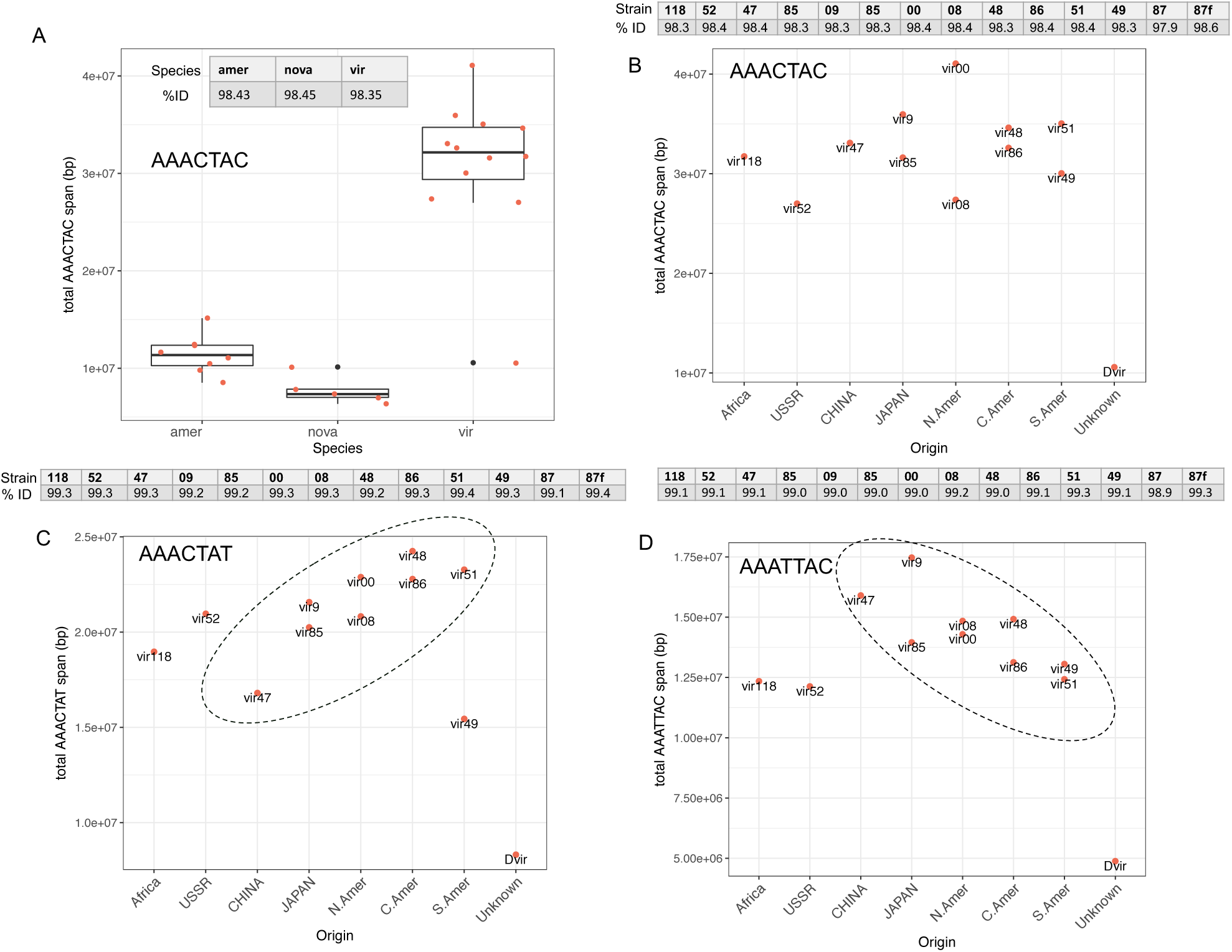
Variation in satellites across species and strains. (**A**) AAACTAC total abundance across the three species. (**B**) AAACTAC, (**C**) AAACTAC, and (**D**) AAATTAC, abundance across *D. virilis* strains originating from different localities (x axis). The strain Dvir is strain 87, the inbred strain used for genome assemblies.

Satellite abundances in *D. virilis* displayed a pattern that appeared to be correlated to the geographic location from which strains were collected. For the centromeric satellite AAATTAC, there was a linear decrease in abundance from West to East then South following probable migration from Beringia (Throckmorton 1982) beginning in China (Figure 4C). For the centromeric satellite AAACTAT, the pattern was the opposite; a linear increase in abundance from West to East then South (Figure 4D). We also analyzed sequence variation in the satellite repeats across strains and species to determine if there were any interesting patterns. On average, the centromeric satellite arrays were very homogeneous (average above 99% sequence identity in Illumina reads). However, AAACTAT had a slightly higher sequence identity than AAATTAC (Figure 4 C and D). There was no pattern in average sequence identity with respect to geography for any of the satellites (Figure 4 C and D). The pericentromeric satellite AAACTAC has almost identical sequence divergence across the three species (∼98.5%). When comparing between a male and female of the same strain, the male had lower sequence identity. In *D. americana* and *D. novamexicana*, the AAACAAC satellite had lower average sequence identity at 97.5%.

There were also several other satellites besides the most abundant four that varied between the *D. virilis* group species. AAACAAT was either absent or in low abundance in *D. virilis* and *novamexicana*, but present between 100,000 - 250,000 copies in *D. americana,* indicating a very recent expansion of this satellite. AAAAAC was present in ∼100,000 copies in *D. americana* and *D. novamexicana*, whereas it is almost absent in *D. virilis*. To differentiate Y-specific satellites, we included a female of strain 87, the reference genome strain, in our Illumina sequencing run in addition to a male. There were several Y-specific satellites in *D. virilis* found by Wei et al. 2018 and validated by our data, which varied between species in the group (AATAATAG, AATAGATT, and ACATAT). All had different patterns of relative abundance between species (https://github.com/jmf422/D_virilis_satellites/blob/master/Intra_inter_species_sequencing/virilis_group_intra-inter_species.html). There was no detectable difference in centromeric or pericentromeric satellite abundance in *D. americana* strains with vs. without the polymorphic centromere-centromere fusion. We conclude that molecular events surrounding the fusion did not produce any changes in satellite abundance (Figure S8).

Amplified repeats of the DINE-1 helitron have been found on the *D. virilis* 5th and Y chromosomes (Dias *et al.* 2015). We also examined variation in this satellite abundance by counting reads that mapped to this family of satellites. There was no striking pattern with respect to geography, however the strain with the highest AAACTAC content had the lowest DINE-1 content, besides the inbred strain 87, which had lower satellite content all-around (Figure S9). We expect AAACTAC and DINE-1 to be both in the pericentromeric region of Chr5 in *D. virilis* based on our FISH results and the previous work. The male contained 2.6x more DINEs than the female of the same strain, indicating that ∼70% of the DINE-1 repeats in the genome are located on the Y chromosome in *D. virilis*.

## Discussion

Here, we used the satellite DNA-rich genome of *D. virilis* to highlight three previously uncharacterized mechanisms for biases that occur in sequencing and analyzing satellite DNA. We emphasize that comparing satellite DNA amounts between different platforms (e.g. Illumina, PacBio, Nanopore, and even different versions of each) should be done with caution as each technology has its own biases. We have found that issues arise when long arrays of simple satellite DNA are attempted to be sequenced by long-read platforms. In the case of PacBio, systematic errors in base calling may be introduced when sequencing through long arrays of satellites. This issue is not specific to our satellites, as a recent study has also found systematic errors and strand biases in shorter arrays of human satellites in both PacBio and Nanopore reads (Mitsuhashi *et al.* 2019). Circular consensus sequencing (CCS) or “HiFi”, a type of sequencing offered by PacBio which allows an accurate consensus to be produced after multiple rounds of sequencing the same molecule, may be more appropriate for sequencing analysis of satellite DNA. No systematic errors in satellite sequences resulted with the new CCS platform after collaboration with PacBio representatives. In the case of Nanopore, it is possible that a similar satellite-specific base calling errors exists, or that there is a strand-specific difference in secondary or tertiary structures that occur in long strands of simple satellite DNA. We caution readers in interpreting simple satellite DNA from long-read sequencing data and suggest validation with satellites of known sequence and abundance if available or Illumina reads (without quality filtering). Long read platforms are already improving their chemistry and software for better satellite characterization. Because long reads are likely to cross boundaries of different repetitive regions, long read sequencing proved useful in understanding the length of the satellite arrays and TE insertions into them. Moreover, we demonstrate that the abundance of satellites in pericentromeric heterochromatin are underestimated when sequencing flies compared to pure diploid tissue because of polyteny. We caution readers in performing quality filtering before simple satellite analysis, as satellite containing reads may be enriched for lower quality scores.

From comparative analysis of satellites, we found the abundant AAACTAC family of satellites arose in the branch leading to the *virilis* phylad 4.5-11 MYA (Figure 2A). Interestingly, the most abundant satellite in *D. montana*, 7-11 MY diverged, is AAAC. The AAACTAC and AAAC satellites were likely derived from a common ancestor satellite (Fig 2A). From both FISH and sequencing analysis, we found that *D. virilis* has the highest total amount of AAACTAC family satellites, *D. novamexicana* has about half of *D. virilis*, and *D. americana* intermediate between the two species. *D. lummei* has the lowest relative satellite content, and its only high-abundance simple satellite is AAACTAC. Unlike the pericentromeric satellite, the centromere-proximal satellite sequence has turned over between *D. virilis* and *D. americana/novamexicana*. The AAACAAC satellite likely evolved in the branch leading to the virilis phylad since it is present in three of the four species studied (Fig 2A). AAACAAC is present in *D. virilis* in relatively low amounts, whereas it became the centromeric satellite on almost all chromosomes in *D. americana* and *novamexicana*. The AAACTAT and AAATTAC satellites are unique to *D. virilis* and occupy the centromeric region. The emerging pattern is that the centromere-proximal satellites have turned over more rapidly than the pericentromeric satellite. This is likely due to satellites participating in conflicts at centromeres (Bayes and Malik 2006, and discussed below). Although sequencing quantified only up to 30 Mb of the AAACTAC family of satellites, FISH confirmed that these satellites are extremely abundant in *D. virilis* and the 40% of the genome estimate seems realistic.

We can make hypotheses about how and why the satellites expanded in *D. virilis*. We know that mutation rates for changes in copy number of satellite DNA are high, and potentially have a tendency to expand rather than contract in the absence of selection (Flynn *et al.* 2017, 2018). High rates of mutation must be accompanied by a regime that would allow a satellite copy number increase to sweep the population - which could be mediated by positive selection if there is a benefit of the satellite increase, or centromere drive if the phenomenon is at play. Alternatively, in a situation where satellites are slightly deleterious, small effective population sizes in isolated populations or continued bottlenecks could allow satellites to expand in the genome without being removed by selection. However, *D. novamexicana* has the lowest effective population size of the virilis phylad and yet it has the lowest amount of satellite DNA. We already know that the centromere-to-centromere fusions in *D. americana* have undergone meiotic drive hypothesized to be mediated by the increase in centromere total size with the fusion (Stewart et al. 2019). The mechanism allowing drive in *D. americana* may have been at play in the branch leading to *D. virilis* or may be currently occurring. Why have satellites not expanded to this extent in the other species? *D. virilis* might have some attributes about its biology that made the satellite expansion favorable or allowable. For example, genome size is positively correlated with development time in Drosophilidae (Gregory and Johnston 2008). *D. virilis* has a slow development time, and this may have evolved in concert with the expansion in satellite abundance in its genome.

We can use data from multiple strains to make hypotheses about factors driving satellite DNA evolution in *D. virilis*. Ancestrally, *D. virilis* had a relatively small effective population size in an isolated range in Asia, and has undergone a recent population and range expansion (Mirol et al. 2008). The amount of the most abundant pericentromeric satellite AAACTAC, does not show a geographical pattern across the global strains. Assuming we sequenced a strain from the ancestral range, this suggests that population bottlenecks were not what allowed AAACTAC to expand, and the satellite expansion likely occurred before the population expansion.

Our observation of rapid evolution and enrichment of AAACTAC in *D. virilis* in a short evolutionary time period (a few million years) is consistent with the centromere-drive model to account for the evolution of centromere complexity in genetic conflict (Malik and Bayes, 2006). In this model, the asymmetric female meiosis can cause competition between the centromeres with or without newly formed satellites or with more or less satellites, to be included into the oocyte to pass to next generation. A consequence of the competition would be runway expansions of centromeric satellites, and rapid replacements by novel satellites. We hypothesize that the pattern of the centromere-proximal satellite AAACTAT increasing on a geographical gradient while AAATTAC decreases along the same gradient is driven by centromeric conflicts. AAACTAT may be starting to occupy centromeres that AAATTAC occupied, benefitting from a transmission advantage (centromere drive), while the AAATTAC satellite may be decreasing in parallel because of selection “pushing back”, for example because of a maximum limit on satellite amount in the centromeric region. Another line of evidence that centromere related conflicts are playing a role is the rapid rate of turnover of the centromere-proximal satellites compared to the pericentromeric satellite.

Interestingly, in *D. novamexicana*, AAACTAC was greatly reduced in the pericentromeric regions on all chromosome pairs except one. Based on the FISH images in *D. novamexicana* and *americana*, we hypothesize that it is the 5th chromosome that has the high amount of AAACTAC satellite. This is interesting because previous work has shown that the 5th chromosome contains a high amount of DINE-1 helitron satellite in *D. virilis* but not in *D. americana* (Dias et al. 2015). This may be evidence of past and ongoing competition and trade-offs between the DINE-1 satellite and AAACTAC. We found that all chromosomes including Chr5 contain a large amount of AAACTAC in *D. virilis*. DINE-1 had a relatively consistent amount across different *D. virilis* strains, however the strain with the highest AAACTAC amount is an outlier with a lower DINE-1 amount (Figure S9). This may indicate a maximum threshold of satellites was reached on this chromosome, and one satellite had to reduce its abundance. We have seen evidence for this trade-off, or appearance of competitive exclusion, being invoked under selection in our previous studies (Flynn *et al.* 2017, 2018). There may have been a similar conflict on Chr5 of *D. novamexicana*, where AAACTAC retained a high copy number to prevent DINE-1 from expanding. Interestingly, the opposite has occurred on the *D. novamexicana* and *D. americana* Y chromosome, where AAACTAC family satellites are absent but DINE repeats are abundant. A potential mechanism mediating apparent stabilizing selection on total satellite abundance is that satellites can act as a sink for heterochromatin factors, with their abundance affecting chromatin state (Lemos *et al*. 2010).

The AAACTAC satellite has remained conserved in sequence and location in the virilis phylad. It has also maintained high levels of sequence identity that is equal in the three species we sequenced (98.5% based on Illumina reads). The conservation may reflect a constraint due to selection or a pervasive mechanism of concerted evolution. The periodicity of the sequence may stabilize the DNA helix wrapping around nucleosomes, or it may be constrained by coevolution of an important satellite DNA binding protein (Maio *et al.* 1977; Jagannathan *et al.* 2018). Additionally, within the AAACTAC family, the position and identity of the four A-nucleotides are conserved in all four satellites (<u>AAA</u>CT<u>A</u>C, <u>AAA</u>TT<u>A</u>C, <u>AAA</u>CT<u>A</u>T, <u>AAA</u>CA<u>A</u>C) - which may indicate constraint based on the above mechanisms. Conservation of particular satellite unit lengths and “AA” periodicities have been found in other divergent species (Lowman and Bina 1990). Concerted evolution of satellites could be achieved by repeated recycling of units by copy number changes associated with replication slippage or unequal recombination or gene conversion (Walsh 1987; Elder and Turner 1995). However, recombination in the pericentromeric heterochromatin has never been detected in wild-type flies (Mehrotra and McKim 2006; Hughes *et al.* 2018). On the other hand, if recombination were occurring, satellite arrays will eventually be lost unless they are conserved by selection (Charlesworth *et al.* 1986). Clearly, we are still lacking in understanding how and why long simple satellite arrays maintain their homogeneity, and whether recombination plays a role in their dynamics. Concordant with the hypothesis that recombination is playing a role, males have lower average sequence identity in the 7 bp satellites than females, which could indicate increased decay on the Y chromosome where there no homologous recombination (Figure 4A).

Moreover, our results suggest that transposable elements flank the AAACTAC satellite array (likely more distally to the centromere) and some TEs have inserted within the array. Our analysis suggests that most TE insertions are recent, but because of the under-representation of satellite containing reads, we cannot estimate the number of TE insertions that have occurred into the satellite arrays. Our analysis is also not precise enough to determine if there are islands of TEs at the centromere in *D. virilis* as has been demonstrated in *D. melanogaster* (Chang et al. 2019).

We found no difference in centromeric and pericentromeric satellites abundances between *D. americana* strains that differ in their X-4 fusion status. This suggests that the fusion event did not result in a large loss of satellites, making the total centromere and pericentromere size is indeed larger on the X-4 fused chromosome than the single unfused chromosome, concordant with the hypothesis that a larger centromeric region results in centromere drive (Stewart et al. 2019). Another interesting observation from the sequencing of multiple strains of the three species was that in all cases, the inbred strain that the reference genome was made from had the lowest amount of AAACTAC. For *D. virilis*, this difference was extreme. It is tempting to speculate that the process of inbreeding and/or long periods in the lab may have driven the reduction in pericentromeric satellite abundance.

In conclusion, our results show very rapid dynamics in the abundant satellites of the *D. virilis* group that are likely explained by various cellular and population-level forces that are not yet understood. Further studies can test if there is a species-specific upper limit to satellite amount per genome or per chromosome upon which negative fitness effects occur, which may result in trade-offs or competition between satellites. Centromere drive may be an important process affecting satellite evolution in this species group, and might partially explain why the satellites expanded 4.5-11 MYA, why satellite sequences at the centromere turned over more rapidly, and why there is a gradient of increasing satellite content related to geographical distribution of strains. A more extensive study to determine if inbreeding or extended periods in the lab drives a reduction in satellite abundance will help illuminate the processes that are important for maintaining satellite content. Determining the frequency of recombination in the large pericentromeric heterochromatin blocks in species like *D. virilis* will be challenging but important for understanding how the satellites maintain homogeneity in their sequence. To understand the role of satellites and the importance of their sequence, unit length, and abundance, researchers can strive to develop methods to engineer satellites by modifying specific bases and their abundances.

## MATERIALS AND METHODS

All scripts for analyzing the data and to produce the results we show are here: https://github.com/jmf422/D_virilis_satellites. Illumina sequencing reads generated for this study are deposited in NCBI SRA under accession PRJNA548201. Raw PacBio reads will be deposited under the same accession. Both will be released upon publication.

### Characterizing satellite DNA from genome assemblies

All scripts and R markdown files used for this analysis are provided in https://github.com/jmf422/D_virilis_satellites/tree/master/Genome_assembly_analysis. We used genome assemblies produced by the PacBio sequencing project (https://www.ncbi.nlm.nih.gov/bioproject/?term=txid7214[Organism:noexp]) of *D. virilis*, *D. novamexicana*, and *D. americana*. We also downloaded the *D. virilis* genome produced by Nanopore sequencing from (Miller *et al.* 2018), and the CAF1 assembly from (Drosophila 12 Genomes Consortium *et al.* 2007). We used Phobos (https://www.ruhr-uni-bochum.de/spezzoo/cm/cm_phobos.htm) and Tandem repeats finder (Benson 1999) to characterize simple and complex satellites in these genome assemblies. To identify the chromosomal linkage of complex satellites in the genome assembly, we produced a dotplot with D-GENIES (Cabanettes and Klopp 2018).

### Characterizing satellite DNA from raw long reads

Characterizing and quantifying satellites from long reads is a challenge because of the sequencing high error rate. We used two approaches to characterize satellites from raw long reads. The first approach, we call k-Seek + Phobos, in which we first broke the reads into 100 bp subreads and ran k-Seek on them. k-Seek is very efficient for analyzing many reads, however is not very sensitive for reads with a high error rate since it was designed for Illumina reads (Wei *et al.* 2014). If k-Seek found satellites on at least one subread, we would run the complete parent read through Phobos. Phobos is more sensitive to imperfect repeats and error rates, but cannot handle huge quantities of data; thus why we only ran the portion of reads identified by k-Seek to have tandem repeats. This approach allowed us to characterize satellites *de novo* and quantify them. All scripts for the analysis of long reads with the k-Seek + Phobos approach are located here: https://github.com/jmf422/D_virilis_satellites/tree/master/LongRead_kseek_Phobos. The second approach we used is Noise-Cancelling Repeat Finder (NCRF, (Harris *et al.* 2019)). This program was designed to quantify satellites from long reads with high error rates. However, it cannot identify satellites *de novo* and requires specific satellite sequences to search for. NCRF also requires a “max divergence allowed” parameter, which we tuned with simulations (see below). Scripts used for the NCRF approach are located here: https://github.com/jmf422/D_virilis_satellites/tree/master/LongRead_NCRF.

We did simulations to assess both approaches: https://github.com/jmf422/D_virilis_satellites/tree/master/Simulations. First, we created a simplified mock *D. virilis* genome with a satellite DNA composition estimated from our FISH results. We could not use the genome assembly because it contained very little satellite DNA. Specifically, each chromosome had a centromeric satellite either AAATTAC or AAACTAC followed by the pericentromeric satellite AAACTAC, combined taking up 40% of the genome. The non-satellite DNA portion of the genome was generated randomly with a 40% GC content. We then used PBSim (Ono *et al.* 2013) to simulate PacBio reads and we used these simulated reads for multiple analyses. First, we used them to tune the max divergence parameter of NCRF by running NCRF repeatedly with max divergence parameters ranging 18-30%. We found that the amount of satellites found, particularly the most abundant one, levelled off at 25% max divergence. This is the parameter value we used moving forward. We also used these simulated reads to quantify satellites with both approaches and compare them. Finally, we used these simulated reads to assess strand biases in long read sequencing data (see below).

### Identification of biases in simple satellites in long read data

We suspected that there were biases in the satellite DNA found in the *D. virilis* group PacBio (and Nanopore) data because we found high abundance satellites that had never been found before with other types of data, and so we suspected they were artifactual. These artifactual satellites were found with both kSeek + Phobos and NCRF approaches, but were not found in the simulated data. We tried testing for a strand bias in reads that contained satellite DNA. Using both the summarized output from NCRF and validated with a custom script (LongRead_NCRF/which_strand_pacbio_script.sh), we counted the satellite DNA stretches that originated from each the positive and negative strand. The positive strand is defined as the one that contains the satellite AAACTAC and derivatives (more As than Ts), and the negative strand is the one that contains the reverse complement (e.g. GTAGTTT, more Ts than As). We did this for the three satellites used in the simulated data and real and artefactual satellites found in the PacBio and Nanopore data. Detailed analysis and visualization of the biases is shown here: LongRead_kseek_Phobos/longread_analysis.html

### Sequencing of polytene and non-polytene tissue

To acquire *D. virilis* pure diploid tissue, we dissected male 3^rd^ instar larvae and collected imaginal discs including the eye-antennal disc and wing discs. Approximately 100 larvae were required to get enough DNA (>1 ug). We also collected ∼5 adult flies for fly libraries. We used the inbred genome assembly strain 87 for these libraries. DNA was extracted with Qiagen DNeasy blood and tissue kit and PCR-free libraries were prepared. Libraries were run on an Illumina NextSeq with 1 x 150 bp reads, and each sample took up approximately 7% of the flowcell. The other libraries run on this flowcell were from an unrelated project including RNAseq from other species. Reads were analyzed with k-Seek both before and after filtering with Trimmomatic (Bolger et al. 2014). FastQC was run to evaluate the quality of the reads. Scripts are here: https://github.com/jmf422/D_virilis_satellites/tree/master/Polyteny. We also analyzed publicly available *D. melanogaster* data from the same strain and same sequencing platform of embryos (non-polytene), salivary glands (extreme polyteny) from (Yarosh and Spradling 2014), and flies (varied levels of polyteny) from (Gutzwiller *et al.* 2015).

### Fluorescence in situ hybridization of satellite DNAs

We followed the protocol of (Larracuente and Ferree 2015) for satellite DNA FISH. We ordered the following probes from IDT with 5’ modifications: (AAACTAC)_6_ with alexa-488 fluorophore, (AAACTAT)_6_ with Cyanine5 fluorophore, (AAATTAC)_6_ with Cyanine3 fluorophore, (AAACAAC)_6_ with Cyanine3 fluorophore, and (AAACGAC)_6_ with Cyanine5 fluorophore. We hybridized three probes at a time, to allow for similar probes to compete to result in specific hybridization with the rationale shown in (Beliveau *et al.* 2015). Hybridization temperature was 32°C. We imaged on an Olympus fluorescent microscope and Metamorph capture system at the Cornell Imaging Facility. Composite images were produced with ImageJ.

### Characterizing TEs linked to satellites and satellites anchored to the genome assembly

We extracted the reads identified to have the 7 bp satellites on them from NCRF results, and then we ran RepeatMasker (http://www.repeatmasker.org) on these reads using parameters: “-nolow” and “-species Drosophila”. All reads had at least 500 bp of tandem satellite on them according to NCRF default parameters, and to avoid spurious identification of TEs from semi-repetitive fragments, we described only TEs in reads that also had at least 500 bp of a TE identified from RepeatMasker. We also BLASTed the same satellite reads to the genome assembly to evaluate if satellite reads could be anchored to the genome assembly. Analysis scripts are here: https://github.com/jmf422/D_virilis_satellites/tree/master/TEs_satellites.

### Sequencing of multiple *D. virilis* group strains

We obtained as many strains of *D. virilis* that have information about where they were collected as possible. This included 12 strains as live stocks we obtained either from stocks in our lab or from the Drosophila species stock center (Table S1). We also prepared a female library for strain 87 for when we wanted to differentiate Y-specific satellite patterns. We also obtained five strains of *D. novamexicana* and eight strains of *D. americana*. All were obtained from live stocks and the inbred genome strains were included for both species as well (strain 14 and G96, respectfully). For *D. americana*, we included four strains that have the chromosome X-4 fusion and four strains that do not have it, based on communication with the Bryant McAllister lab. DNA was extracted as above from five flies each and samples were prepared identically as above and sequenced on 50% of 3 flowcells of Illumina NextSeq 1 x 150 bp reads. We dispersed the samples from each species between multiple flowcells. Our samples took up only half the flowcell with the other half being occupied by a RNAseq libraries from an unrelated project.

All scripts used to analyze these data are located here: https://github.com/jmf422/D_virilis_satellites/tree/master/Intra_inter_species_sequencing. Reads were evaluated with FastQC and not filtered for quality based on the potential bias of Illumina quality scores on satellites. We used k-Seek to quantify satellites. We tried several normalization strategies but decided the most appropriate was a mapping-free normalization. We estimated average depth by dividing the total number of bases sequenced by the estimated genome size by flow cytometry (Bosco *et al.* 2007). We believe this was the best option in this case because: 1) we were concerned about a mapping bias because for each species the strain that the genome assembly was made from may have more reads map to it; 2) after masking the genome from the 7mer satellites and also excluding the X and Y contigs (because we had male and female strains, and the Y chromosome contained more low GC regions) - there was little difference in coverage based on GC content. We include results when we used a mapping based GC normalization in the sub-directory “AlternativeNormalization”.

We used NCRF with modified parameters (minlength=100, maxdiv=10) to characterize the average sequence identity of satellite arrays from the Illumina data. To quantify DINE-1 satellites across *D. virilis* strains, we produced a library of DINE-1 satellite variants based on our PacBio genome analysis. We then mapped Illumina reads to this library and normalized the number of reads that mapped to any sequence in the library by the estimated depth. We also analyzed Illumina DNA sequencing reads of *D. montana* (Parker et al. 2018) and *D. lummei* (Ahmed-Braimah *et al.* 2017) with k-Seek to identify the most abundant satellites and whether or not the AAACTAC satellite family was present.

## Acknowledgments

We thank Yasir Ahmed-Braimah for helpful discussions and advice for some analyses. We also thank Bryant McAllister for providing *D. americana* strains along with their fusion status. We are grateful for Elissa Cosgrove’s help with some computational trouble-shooting and Asha Jain’s help in preparing DNA sequencing libraries. We also thank Danny Miller for useful discussions and for providing the raw Nanopore reads from his study. Sarah Kingan, Jane Landolin, and Greg Young from Pacific Biosciences were very helpful in exploring the causes for the artefactual repeats and in producing the HiFi data. We thank Amanda Larracuente for advice on FISH protocols and the Cornell Imaging Facility for use of their microscope. This project was funded by NIH grant number: GM116113 to RAW, ML, and AGC and GM119125 to AGC and Daniel Barbash. JMF was supported by an NSERC PGS D fellowship.

