## Supplemental for "Evolutionary dynamics of abundant 7 bp satellites in the genome of *Drosophila virilis*"

**Table S1**. Metadata for strains sequenced in this study. D. virilis and novamexicana strains were obtained from the Drosophila Species Stock Center and D. americana strains were obtained from Bryant McAllister's lab.

| Strain | Species | Origin | Notes |
|---|---|---|---|
| vir51 | D. virilis | Chile | |
| vir52 | D. virilis | USSR | |
| vir86 | D. virilis | Mexico | |
| vir47 | D. virilis | China | |
| vir49 | D. virilis | Argentina | |
| vir85 | D. virilis | Japan | |
| vir08 | D. virilis | California, USA | |
| vir00 | D. virilis | California, USA | |
| vir118 | D. virilis | Rwanda, Africa | |
| vir48 | D. virilis | Mexico | |
| vir87 | D. virilis | Unknown | inbred genome strain |
| vir9 | D. virilis | Japan | |
| amMK1012 | D. americana | Indiana, USA | 100% X-4 fusion |
| amCI0518 | D. americana | Louisiana, USA | 0% X-4 fusion |
| amCI0515 | D. americana | Louisiana, USA | 0% X-4 fusion |
| amG96 | D. americana | Indiana, USA | 100% X-4 fusion, inbred genome strain |
| amCI0538 | D. americana | Louisiana, USA | 0% X-4 fusion |
| amMK0738 | D. americana | Indiana, USA | 100% X-4 fusion |
| amSB | D. americana | Iowa, USA | 100% X-4 fusion |
| amML975 | D. americana | Louisiana, USA | 0% X-4 fusion |
| Gnova14 | D. novamexicana | Utah, USA | inbred genome strain |
| nova13 | D. novamexicana | Arizona, USA | |
| nova12 | D. novamexicana | Colorado, USA | |
| nova8 | D. novamexicana | New Mexico, USA | |
| nova4 | D. novamexicana | Utah, USA | |

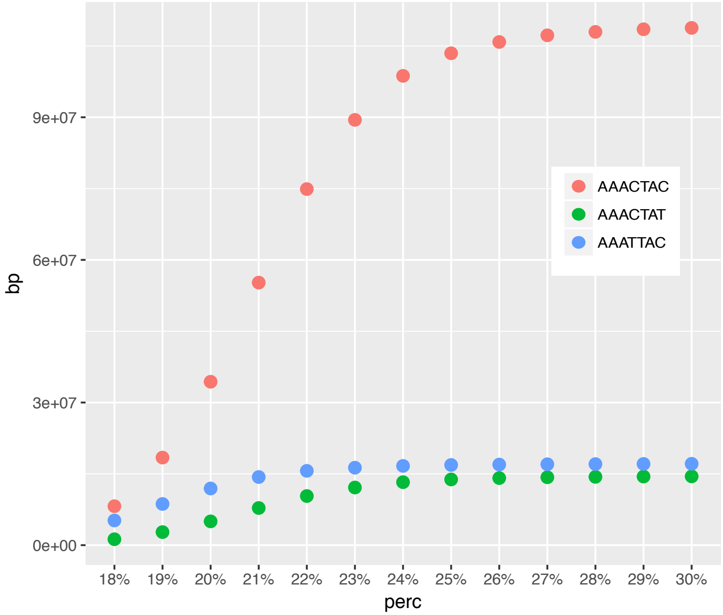


Figure S1. NCRF Simulations results. Running NCRF with different max error allowed on reads from PBSIM off a mock genome composed of perfect repeats..

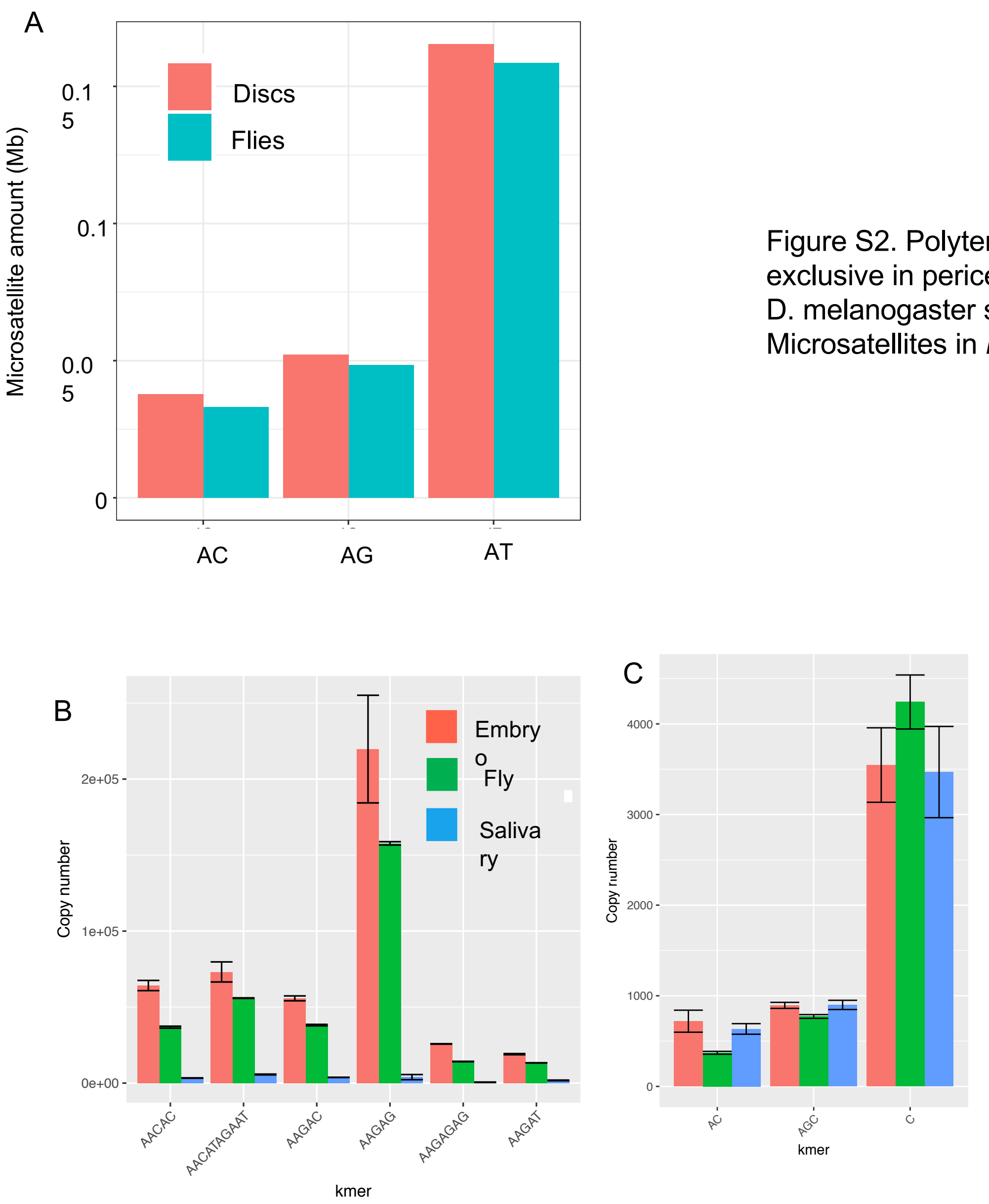


Figure S2. Polyteny vs. diploid analysis. (A) Microsatellites expected to not be exclusive in pericentromeric heterochromatin (control for Fig 2E). (B) Analysis of D. melanogaster sequencing data from embryos, flies, and salivary glands. (C) Microsatellites in *D. melanogaster* data, control for Fig S2B.

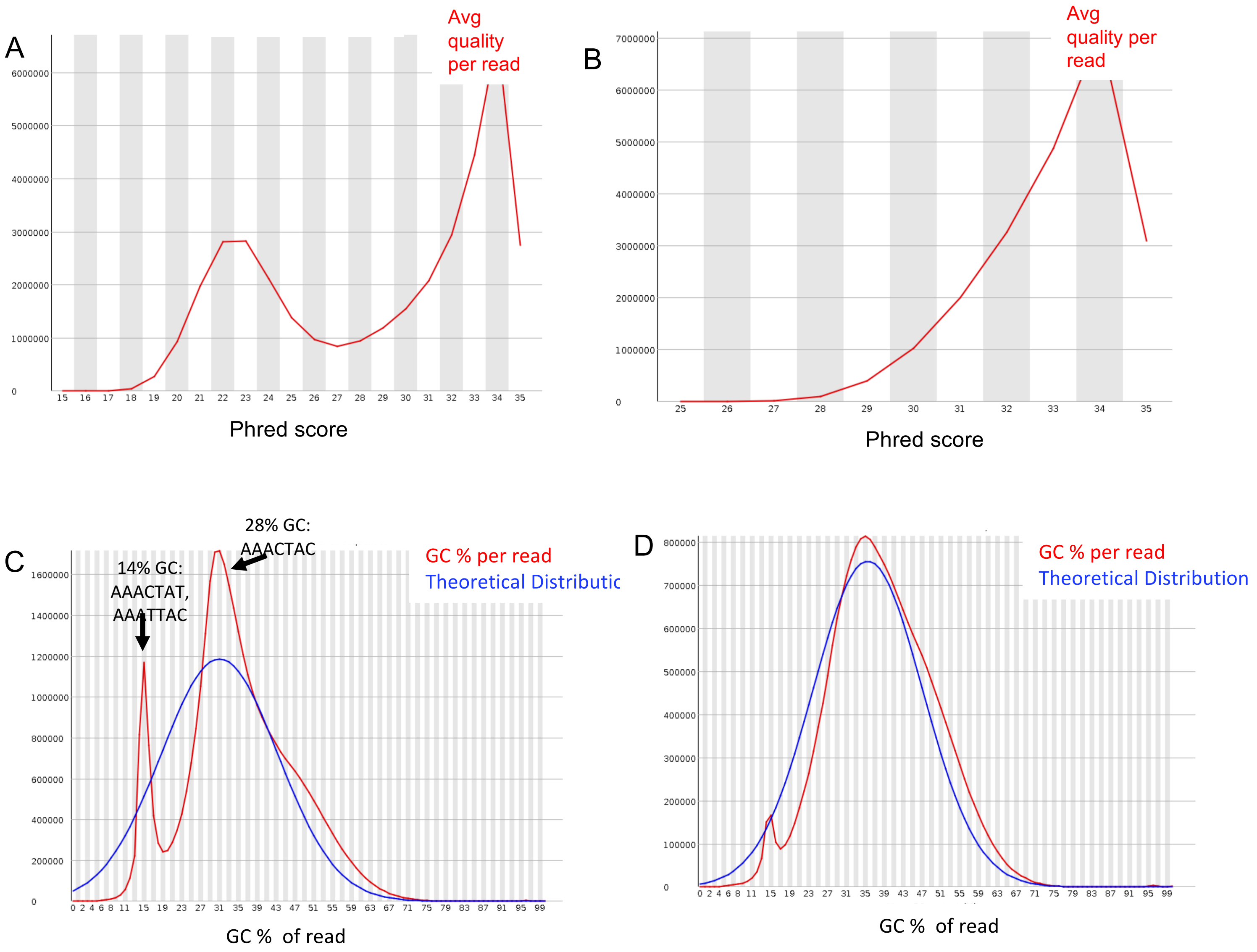


Fig S3. Satellite containing reads are enriched for low quality scores in Illumina data. A) Quality score distribution in the raw reads. (B) Quality score distribution after quality filtering. (C) GC distribution of raw reads. (D) GC distribution of reads after quality filtering.

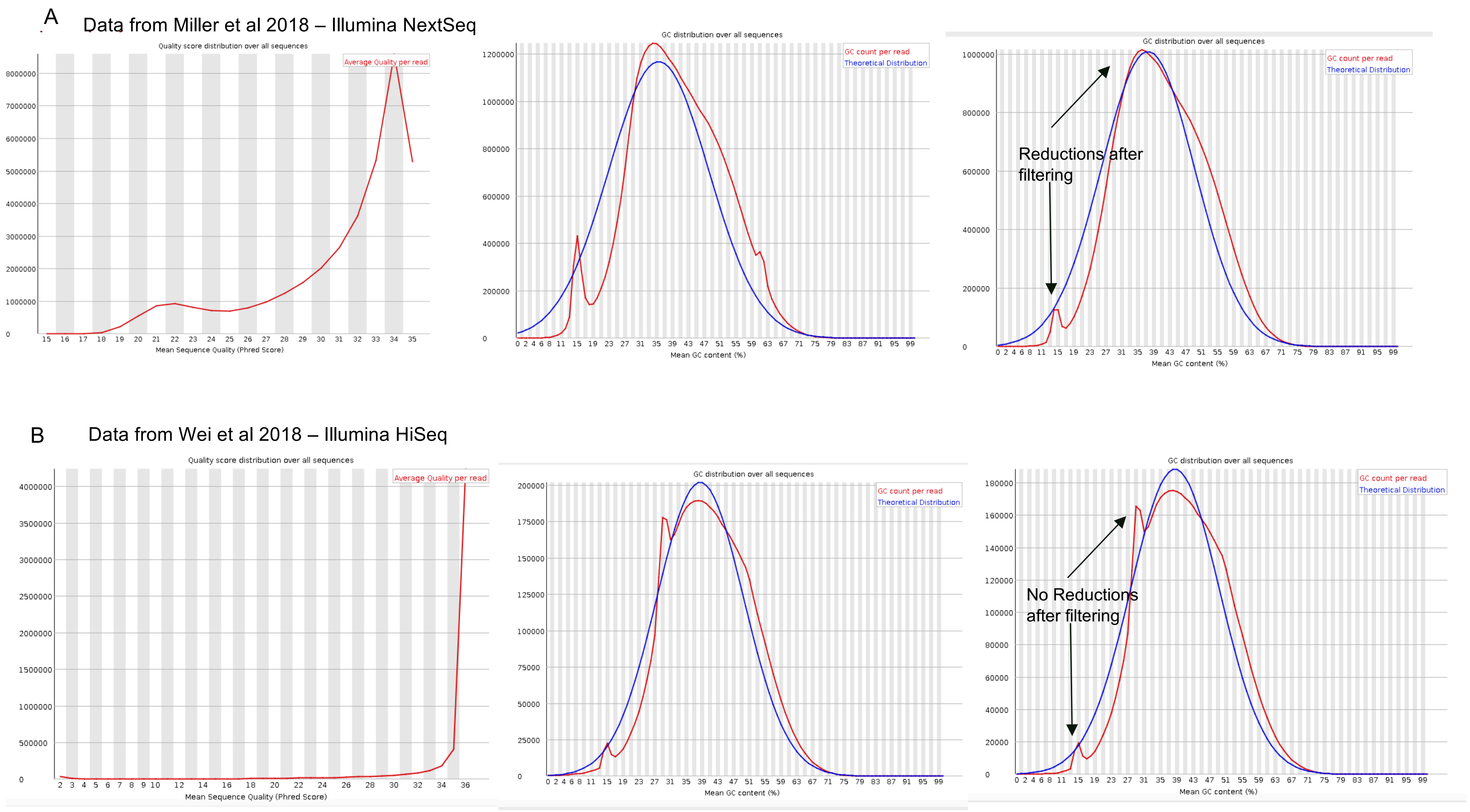


Fig S4. Satellite containing reads are enriched for low quality reads in one Illumina dataset (A) produced on a NextSeq, but not on a second dataset (B) produced on a HiSeq.

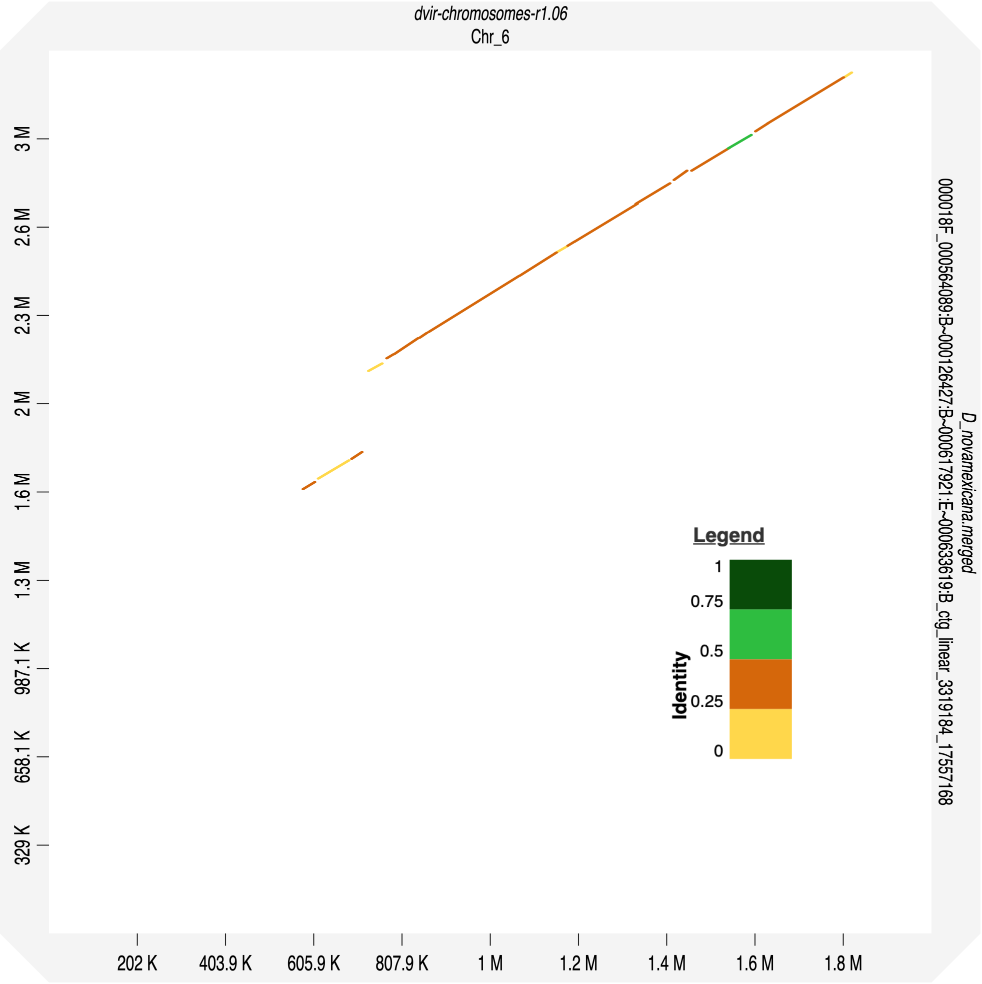


Figure S5. Dotplot of D. novamexicana contig 000018F, which contained a high amount of the 32 bp satellite (from 0-1.5 M) and its similarity to a large region on *D. virilis* chromosome 6.

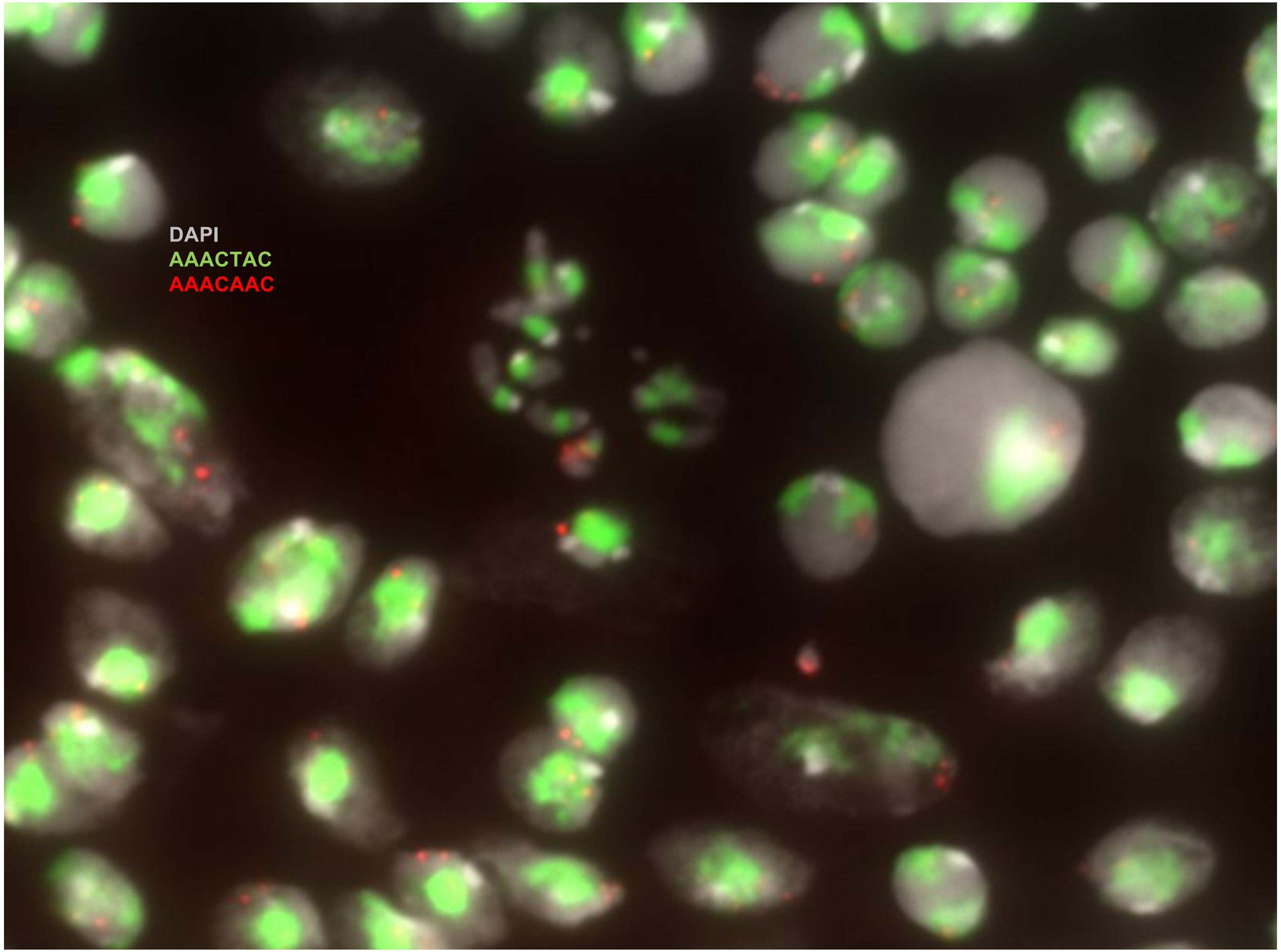


Fig S6. FISH image of D. virilis male. The AAACAAC satellite appears on a single chromosome pair. This is clear from the metaphase chromosomes (middle) as well as the interphase cells where two distinct foci are localized. The distinct foci in interphase cells is in contrast to the AAACTAC satellite, which takes up a large portion of the nucleus.

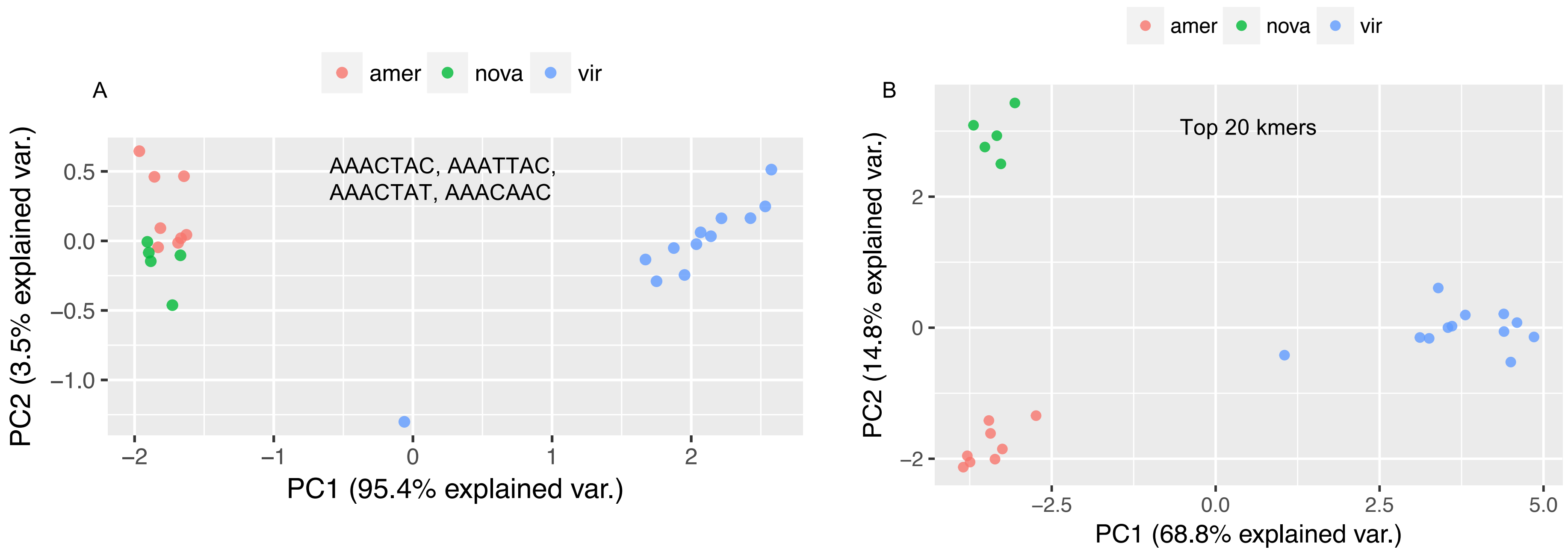


Fig S7. PCA of satellite DNA copy number of D. virilis group strains. (A) Using only the abundances of the satellites AAACTAC, AAATTAC, AAACTAT, AAACAAC. (B) Using the abundances of the 20 most abundant simple satellites.

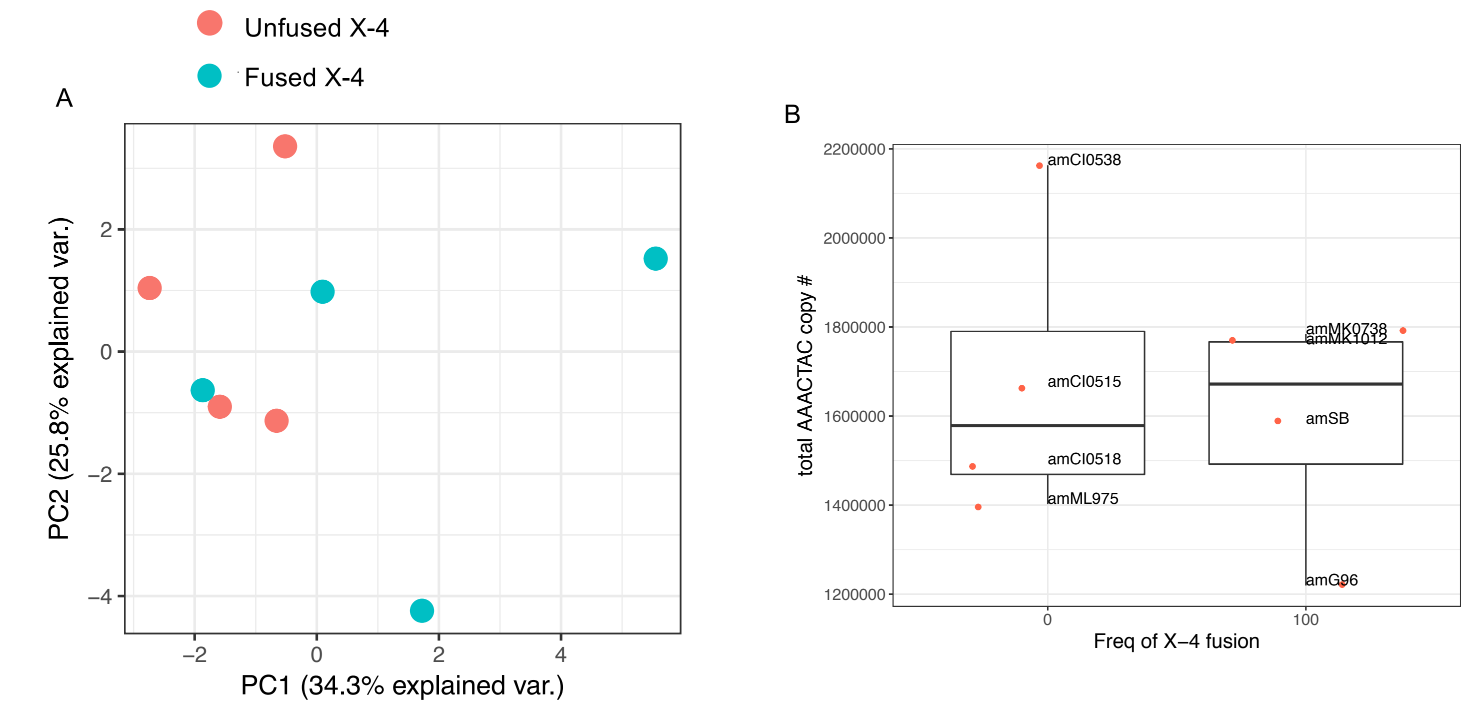


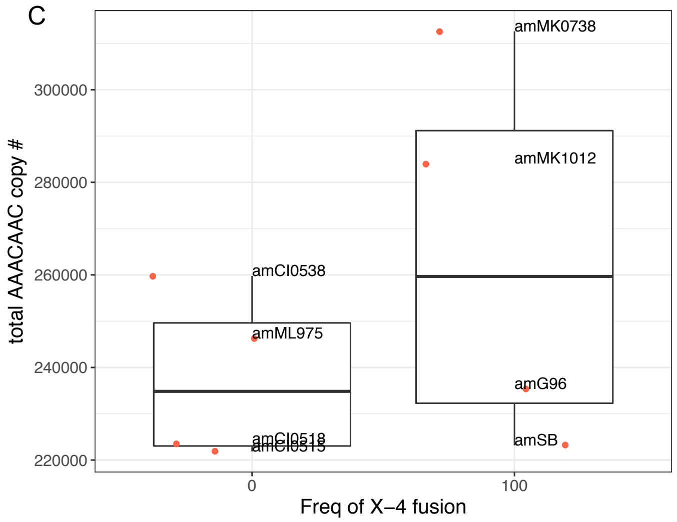


Fig S8. Comparison of D. americana strains with and without the X-4 chromosomal fusion. (A) PCA using the top 20 simple satellites. (B) Boxplot of AAACTAC copy number. (C) Boxplot of AAACAAC copy number.

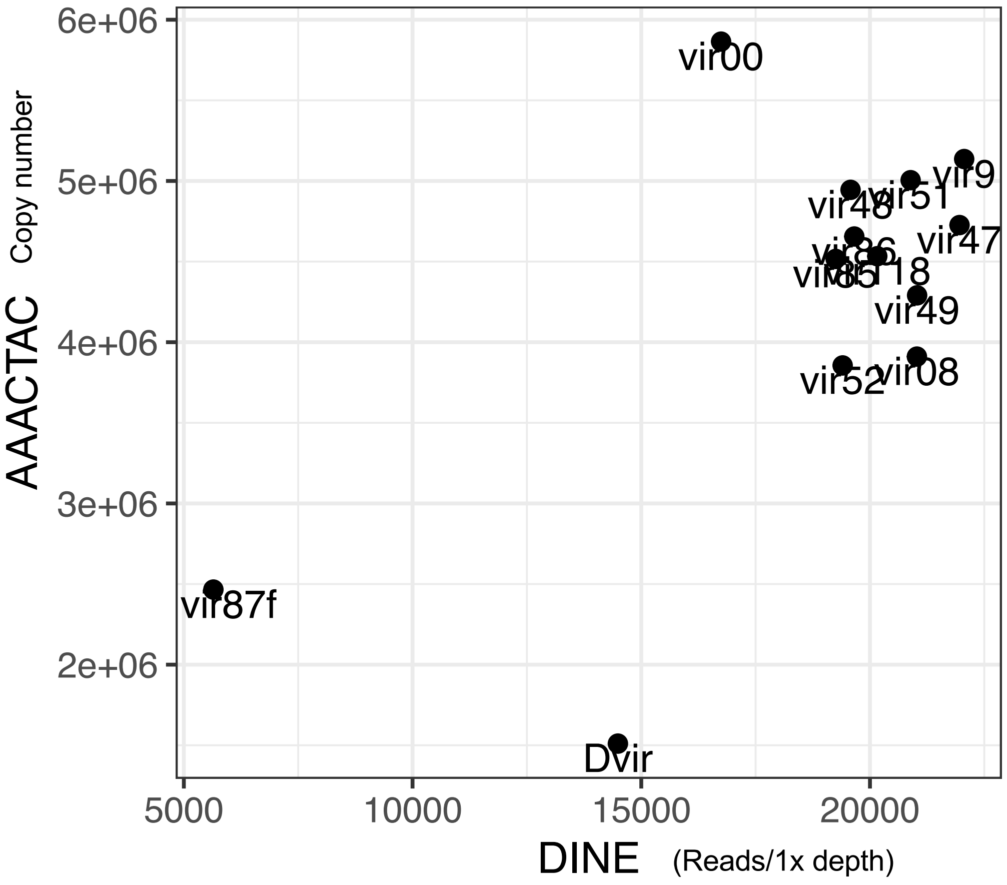


Fig S9. DINE amount vs. AAACTAC copy number. DINE amount is the number of reads mapped to the DINE repeat / estimated depth.
